# Improving *de novo* Protein Binder Design with Deep Learning

**DOI:** 10.1101/2022.06.15.495993

**Authors:** Nathaniel Bennett, Brian Coventry, Inna Goreshnik, Buwei Huang, Aza Allen, Dionne Vafeados, Ying Po Peng, Justas Dauparas, Minkyung Baek, Lance Stewart, Frank DiMaio, Steven De Munck, Savvas N. Savvides, David Baker

**Affiliations:** Department of Biochemistry, University of Washington, Seattle, WA, USA; Institute for Protein Design, University of Washington, Seattle, WA, USA; Molecular Engineering Graduate Program, University of Washington, Seattle, WA, USA; Department of Bioengineering, University of Washington, Seattle, WA, USA; VIB-UGent Center for Inflammation Research, Ghent, Belgium; Unit for Structural Biology, Department of Biochemistry and Microbiology, Ghent University, Ghent, Belgium; Howard Hughes Medical Institute, University of Washington, Seattle, WA, USA

## Abstract

We explore the improvement of energy-based protein binder design using deep learning. We find that using AlphaFold2 or RoseTTAFold to assess the probability that a designed sequence adopts the designed monomer structure, and the probability that this structure binds the target as designed, increases design success rates nearly 10-fold. We find further that sequence design using ProteinMPNN rather than Rosetta considerably increases computational efficiency.

## Introduction

While most current methods for generating proteins which bind to protein targets of interest involve library selection methods or immunization, there has been recent progress in computational design approaches which, in principle, could provide much faster routes to affinity reagents having desired biophysical properties that target specific surface patches. A general Rosetta-based approach to designing binding proteins using only the structure of the target was recently developed and used to design binding proteins to 13 different target sites^1^. Given a specified region on a target of interest, the method designs sequences predicted to fold up into protein structures that have shape and chemical complementarity to the region.

While providing for the first time a general computational route to designing binders to arbitrary protein targets, only a small fraction of designs typically have sufficiently high affinity for experimental detection. There are two primary failure modes (Fig 1a): first, the designed sequence may not fold to the intended monomer structure (Fig 1b), and second, the designed monomer structure may not actually bind the target (Fig 1c). Rosetta frames both the folding and binding problems in energetic terms; the designed sequence must have as its lowest energy state in isolation the designed monomer structure, and the complex between this designed monomer structure and the target must have sufficiently low energy to drive formation of the design-target protein complex. The primary challenges in accurate design of both the monomer structure and the protein-protein interface are inaccuracies of the energy function, and the very large size of the space which must be sampled: if the energy function is inaccurate, or conformational sampling is incomplete, the designed sequence may not fold to the intended monomer structure and/or the monomer may not bind to the target as intended. We set out to investigate whether recently developed deep learning methods could help overcome these problems.

**Figure 1.**
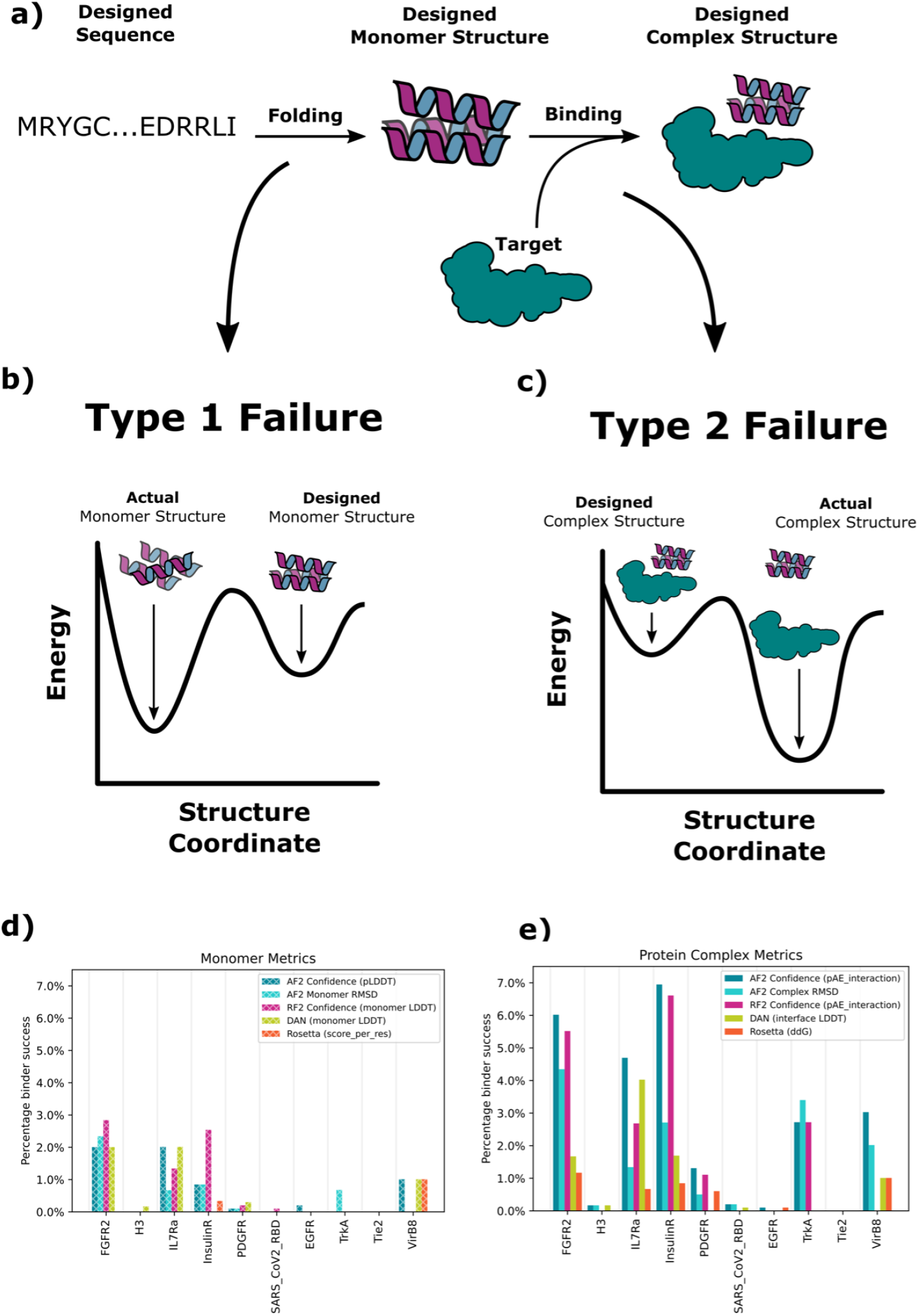
Monomer and protein complex structure prediction metrics distinguish previously designed binders from non-binders. (A) For binder design to be successful, the designed sequence must fold to the designed binder monomer structure (left), and this structure must form the designed interface with the target protein (right). (B-C) Design failure modes. (B) Type-1 Failures. The designed sequence does not fold to the designed monomer structure. (C) Type-2 Failures. The designed sequence folds to the designed monomer structure. (D, E) The retrospective experimental success rate (YSD SC_50_ < 4 μM) for the top 1% of designs selected according to different monomer (D) or protein complex (E) based metrics over 10 targets from Cao et al.

## Retrospective Analysis

We began by investigating the ability of deep learning methods to discriminate binders from non-binders (filtering) in the set of ~1 million experimentally characterized designs for 13 different targets described in Cao et al. 15,000-100,000 designs were experimentally tested for each target, and the number of actual binders ranged from 1 to 584.

### Type 1 failures

We first focused on identifying Type 1 failures (Fig 1) in which the designed sequence does not fold to the intended monomer structure. As a baseline, we used the Rosetta energy of the monomer. Not surprisingly, as this metric was already used as a stringent filter in generating the input scaffold set for the Rosetta interface design calculations^2^, it provided little discriminatory power (Fig 1d). In contrast, the deep learning based accuracy prediction method DeepAccuracyNet^3^ (DAN) was able to partially discriminate binders from non-binders (Fig 1d).

We used AlphaFold2^4^ (AF2) and an updated version of RoseTTAFold^5^ (RF2) to directly assess Type 1 failures by comparing the predicted structure for each designed sequence to the corresponding design model (predictions for the designed monomers were very close to crystal structures where available; Fig S1). We found that the closer the predicted binder structure was to the Rosetta-designed structure in Cα RMSD, and the higher the prediction confidence (pLDDT; Fig 1d), the more likely a binder was to be successful (Fig S4; the pLDDT of AF2 and RF2 were equally discriminative). These results suggest that Type 1 failures contribute to the low success rate of binder design, and that such failures can in part be identified by discrepancies between design models and AF2 or RF2 structure predictions.

### Type 2 failures

To estimate the likelihood of the designed binder structure forming an interface with the intended target, Cao et al. primarily used the difference in energy of the bound complex and the unbound monomers allowing sidechain repacking as computed by Rosetta (Rosetta ddg). Despite its extensive use during the original calculations, Rosetta ddg remains an effective filter (Fig 1e). The DAN complex accuracy metric was similarly effective (Fig 1e).

We next evaluated the ability of AF2 and RF2 to reproduce the five minibinder crystal/cryoEM structures from Cao et al. Given an MSA for the target protein and the single sequence of the designed binder, AF2 predicted the complex structure with binder Cα accuracy between 1.0Å-2.0Å for three of five. To enable AF2 to be used for binding prediction in cases where the target is incorrectly modeled, we initialized the AF2 pair representation with an encoding of the Rosetta binder structure and provided the target structure as a template; this “AF2 initial guess” protocol (see Supplemental Information) was able to recapitulate the crystal structures for all 5 crystallized minibinder interfaces with binder Cα accuracy between 1.0Å-2.0Å RMSD (Fig S1; using a templated target but no initial guess, RF2 recapitulates 4 of 5 between 1.0Å-2.0Å RMSD).

We used the AF2 initial guess approach and RF2 with a templated target but without an initial guess to directly evaluate Type 2 failures by comparing the predicted and designed structures for each complex in the dataset. Designs for which the Cα RMSD of the predicted complex to the design model and the predicted inter-chain error (pAE_interaction) was relatively low, had nearly ten fold higher experimental success rates than the overall population (Fig 1e). Confident predictions had very high success frequencies (see the Receiver Operator Characteristic (ROC) curves in Fig S6) with sharp increases in success rates for designs with pAE_interaction < 10. AF2 had slightly better performance than RF2 (Fig 1e), and we used this in the new design campaigns described in the following section.

## Prospective Analysis

The retrospective analysis in Fig 1 suggests incorporation of AF2 or RF2 into the design pipeline as a final evaluation filter could considerably increase the design success rate. To directly test this hypothesis, we carried out binder design campaigns on four new targets of considerable biological importance: ALK^6^, LTK^6^, IL10 receptor-α^7^, and IL2 receptor-α^8–11^ (two sites). Using the protocol of Cao et al., we generated computational libraries of ~2 million designs for each target and filtered these down to ~20,000 designs to be experimentally tested for each target: ~15,000 designs using the physically-based filters of Cao et al. and ~5,000 designs with AF2 pAE_interaction < 10 (these designs were also filtered by additional metrics as described in the Supplement). Synthetic genes were obtained for the ~80,000 designs, transformed into yeast, and the resulting library sorted for display of the proteins on yeast cells, followed by sorts at 1uM target with avidity, and sorts at decreasing concentrations of target. The frequency of each design at each sort was determined by deep sequencing, and SC50 values (the concentration where half of the expressing yeast-cells are collected) estimated as described in Cao et al. Designs with SC_50_ values better than 4 μM were considered successes; the number of successes for the four targets ranged from 1 to 17. For all four targets, there was a considerably higher success rate (number of successes / number of designs tested) in the AF2 filtered design set than in the Rosetta set (Fig 2b). Physically-based filtering yielded successful binders for two targets: LTK and Site 1 of IL-2Rα; for these the AF2 filtered libraries had 8- and 30-fold higher success rates, respectively. AF2 filtered libraries also yielded successful binders to both ALK and IL-10Rα; physically-based filtering yielded no successful binders to either of these targets (Neither filtering method was able to generate successful binders to Site 2 on IL-2Rα).

**Figure 2.**
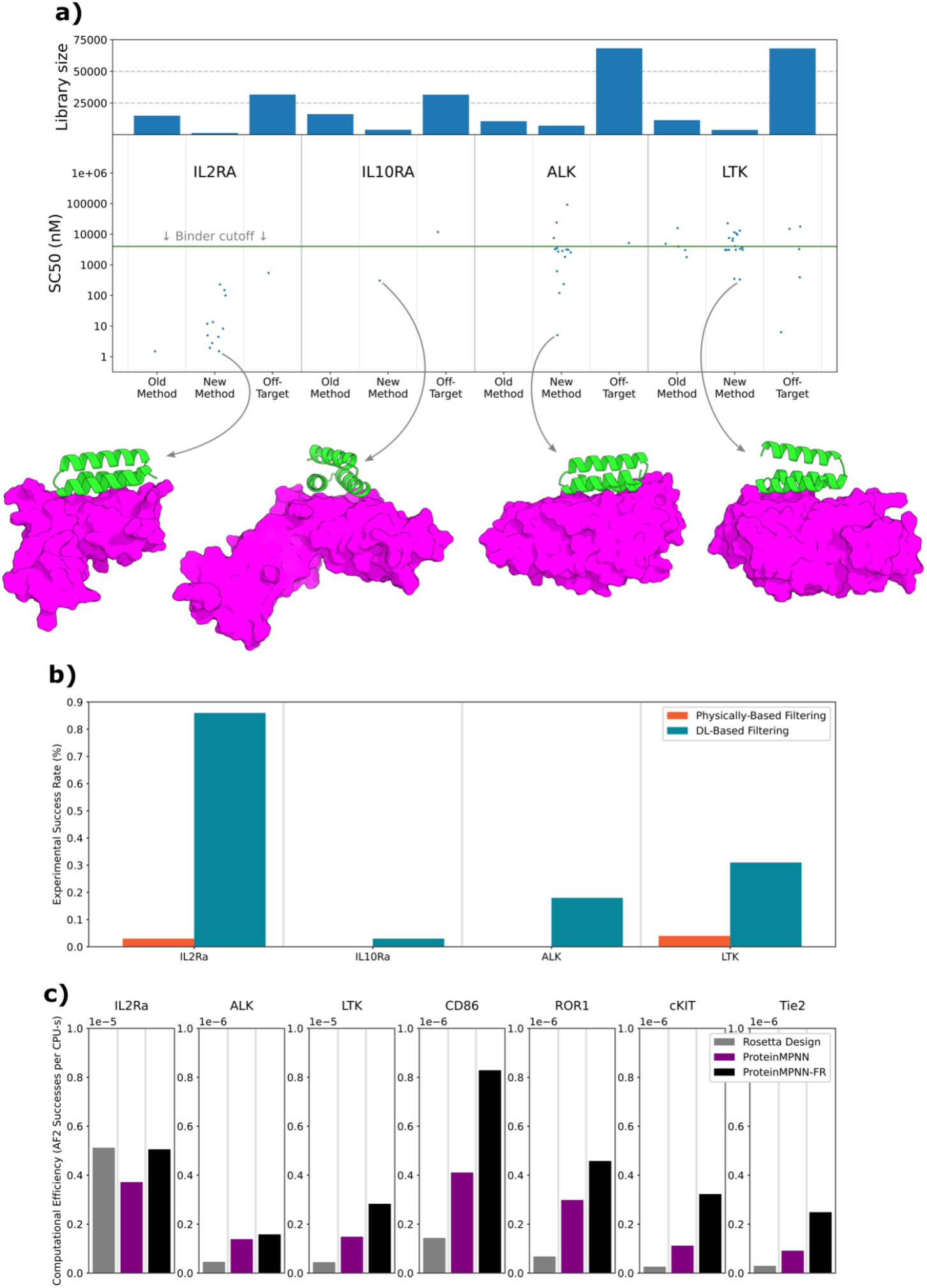
Incorporation of structure prediction metrics increases design success rate on new targets. (A) Results of Prospective Campaigns. For each target the SC_50_ from YSD is shown for all designs which showed binding by YSD (like Kd’s, lower values are better). The number of designs included in each library for each target is indicated by the bars in the top panel. The AF2-predicted structure of the top scoring on-target design is shown as a cartoon. No binders were identified to Site 2 of IL2 receptor-α so this campaign is not included here. (B) Experimental success rate for libraries filtered by DL-based filtering versus Physically-based filtering for the four prospective targets. (C) Computational efficiency (the number of designs with pAE_interaction < 10 per CPU-s) for the original Rosetta sequence design protocol, ProteinMPNN sequence design and ProteinMPNN sequence design plus Rosetta relax over seven targets. Note that IL2Rα and LTK are plotted with a 1×10^-5^ y-axis while the rest of the targets are plotted with a 1×10^-6^ y-axis.

### Increasing binder design pipeline compute efficiency with ProteinMPNN

While an effective predictor of binder success, the AF2 filter is computationally expensive (~30 GPU-seconds per design) and, on average, only ~2.6% of designs pass, so large numbers of prediction calculations must be run. As a metric of computational efficiency, we use the number of designs passing the AF2 filter a method can generate per unit compute time (with a conversion factor of 100 CPU-s to 1 GPU-s because of the relative scarcity of GPU resources):

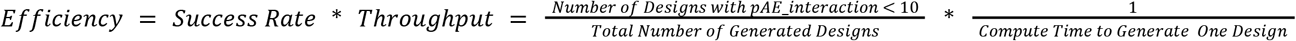

Rosetta-design has an average efficiency of about 8.4×10^-7^ successful designs per CPU-s equivalent (the efficiency of a design protocol is target dependent, per-target efficiencies are shown in Fig 2c).

We investigated whether the recently developed deep learning based sequence design method ProteinMPNN^12^ could be used to increase the efficiency of the design pipeline. ProteinMPNN is very fast, generating a sequence for a minibinder backbone in ~2 CPU-s compared to ~350 CPU-s for Rosetta-design. We first compared the experimental success rate of ProteinMPNN designs to Rosetta designs. For all Rosetta designs from the above prospective study with low complex Cα RMSD AF2 predictions, we generated new sequences using either ProteinMPNN or Rosetta-Design (~10^4^ designs in total). Genes encoding designs with AF2 pAE_interaction < 10 (~10^3^ per method) were synthesized, and the binding evaluated by FACS followed by deep sequencing as described above. We found that the design success rate of ProteinMPNN and Rosetta-design were similar (Fig S7), thus the considerable increase in speed comes with no decrease in performance.

ProteinMPNN design alone had an average efficiency of 8.9×10^-7^ successful designs per CPU-s equivalent. When calculating the fold efficiency improvement for each target and taking the average, ProteinMPNN has ~3-fold higher efficiency compared to Rosetta-design (Fig 2c). Since unlike Rosetta, ProteinMPNN keeps the protein backbone fixed, it is sensitive to the input backbone structure quality. Inspired by the very efficient alternation between sequence optimization and structure refinement in Rosetta flexible backbone design^13^, we evaluated similar cycling between ProteinMPNN and Rosetta structure refinement (FastRelax), hoping to converge on a high-quality backbone that would then allow ProteinMPNN to generate a high-quality sequence. This hybrid ProteinMPNN/Rosetta sequence design protocol (henceforth referred to as ProteinMPNN-FR) generated AF2 pAE_interaction<10 structures at an average rate of ~4.3% with a throughput of 1 design per 120 CPU-s for an average efficiency of 1.4×10^-6^. The average per-target efficiency improvement of ProteinMPNN-FR over Rosetta-design is ~6-fold (Fig 2c).

## Conclusion

Our retrospective and prospective studies suggest an increase in design success rate of nearly ten fold. In contrast to Rosetta energy calculations and DAN structure accuracy measures, which operate on single protein structures (or with Rosetta relax calculations, structures very close to the query), structure prediction calculations implicitly assess the fit of the sequence with the desired target structure compared to all others. As observed previously^14^, such consideration of the overall folding landscape enables considerably more accurate assessment of the likelihood a design will fold and bind as intended compared to evaluation of only the depth of the designed energy well. Although the protocol reported here is an order-of-magnitude improvement over the previous state-of-the-art, it is clear that much about interface energetics remains poorly understood; success rates among the targets remain low (<1%) and no binders were identified to Site 2 of IL2 receptor-α. There is also considerable room for improvement in designing high affinity binders; as with the original pipeline the initially generated binders are in the high nM affinity range. Given the rate of progress in the field, we anticipate further increases in design success rates and affinities in the near future, which will make computational protein design methods even more powerful compared to empirical selection methods for generating affinity reagents and therapeutic candidates.

## Acknowledgments

This work was supported with funds provided by The Donald and Jo Anne Petersen Endowment for Accelerating Advancements in Alzheimer’s Disease Research (N.B.), a gift from Microsoft (J.D., M.B., D.B.), the Audacious Project at the Institute for Protein Design (A.A., D.V., D.B.), a grant from DARPA supporting the Harnessing Enzymatic Activity for Lifesaving Remedies (HEALR) Program (HR001120S0052 contract HR0011-21-2-0012, I.G., F.D., Y.P.P., B.H., L.S., D.B.), the Flanders Institute for Biotechnology (S.N.S), a Strategic Basic Research grant from Research Foundation Flanders (S.N.S.) and the Howard Hughes Medical Institute (B.C., D.B.).

We thank AWS and Microsoft for generous gifts of cloud computing credits.

## Author Contributions

N.B., B.C., and D.B. designed the research; N.B. and B.C. contributed equally; N.B. and B.C. developed the method; N.B. and B.C. designed the binders; I.G., B.H., A.A., Y.P.P., and D.V. performed the yeast screening; J.D. developed ProteinMPNN; M.B. and F.D. developed RoseTTAFold2; S.D.M. and S.N.S. solved the structures of ALK and LTK; all authors analyzed data; L.S. and D.B. supervised research; N.B., B.C., and D.B. wrote the manuscript with input from the other authors. All authors revised the manuscript.

## Data Availability

The raw data from the prospective study presented in Figure 2 is available from the corresponding author upon reasonable request. The Rosetta macromolecular modeling suite (https://www.rosettacommons.org) is freely available to academic and non-commercial users. Commercial licenses for the suite are available via the University of Washington Technology Transfer Office.

## Code Availability

The Rosetta macromolecular modeling suite (https://www.rosettacommons.org) is freely available to academic and non-commercial users. Commercial licenses for the suite are available via the University of Washington Technology Transfer Office. Scripts for running ProteinMPNN-FastRelax and AF2 with templating and initial guess is available from the authors upon reasonable request and will be available with the published peer-reviewed manuscript.

## Supplementary Information

### AF2 Initial Guess

The input structure is first converted to AF2 atom positions using the af2_get_atom_positions function. These positions are then fed, along with the standard model inputs into the AF2 Model Runner. In the AlphaFold class of the AlphaFold code, on the first recycle the prev_pos variable is initialized to the input AF2 atom positions as opposed to the standard initialization of all zeros. A script to run AF2 initial guess and the modified source code is available from the authors upon reasonable request and will be available with the published peer-reviewed manuscript. The AlphaFold model used in the script and in this work is configured to run with a reduced number of extra MSA sequences which speeds the inference of the network dramatically, as described in previous work^15^.

### ProteinMPNN FastRelax

This protocol takes as input a protein complex structure. The ProteinMPNN is then fed the complex structure with the binder sequence masked and asked to assign the binder a sequence. The new sequence is then threaded back onto the binder structure in the complex and the complex structure is relaxed using Rosetta FastRelax. The relaxed complex structure can then be used as the input to the ProteinMPNN to continue the cycle. A python script to perform this design technique is available from the authors upon reasonable request and will be available with the published peer-reviewed manuscript.

### Design and filtering procedure for Prospective Study

The prospective study was performed at a time of rapid protocol discovery with a tight deadline for placing the gene-order. As such, not every experiment that could have been performed was performed. However, the comparison of Rosetta filtering to AF2 filtering was the main goal and the data required for this comparison was plentiful.

The standard procedure from Cao et. al. was followed for the 4 targets starting with the following pdbs: IL2RA (1Z92, 2B5I, 3NFP, 2ERJ), IL10RA (1LQS), ALK (Privately communicated structure. Now 7NWZ), LTK (Privately communicated structure. Now 7NX0). The “recommended_scaffolds.list” from Cao et. al. were used and on the order of 10M RifDock^16^ outputs were generated for each target with about 500K FastDesigned. ~6K motifs were extracted, grafted up to 10M docks, and 500K FastDesigned again. The resulting 1M designs for each target were predicted by AF2.

From this set of 1M designs, 3 overlapping subsets were selected. The first subset was the Rosetta-control group where the AF2 predictions were ignored and the top ~18K per target were selected by the pareto-front method from Cao et. al. looking at target_delta_sap, ddg, contact_patch, and contact_molec_sq5_apap_target. The second subset was the AF2-filtered group where all designs passing pae_interaction < 10 and af2_complex_rmsd < 5Å were included. This set was typically around 8K per target. The third subset was all predictions with af2_complex_rmsd < 5Å. These designs were designated to be redesigned and were typically about 12K in scale.

These AF2-predicted interfaces were then designed either with Rosetta or ProteinMPNN. Here, ProteinMPNN was used to generate a protein sequence from the input coordinates and no further optimization was performed. The Rosetta-redesigned and ProteinMPNN-redesigned pools were predicted again by AF2 and were filtered either with the Rosetta filters mentioned above or the AF2 filters mentioned above resulting in pools of sizes 9K (Rosetta-Rosetta), 2K (Rosetta-AF2), and 2K (ProteinMPNN-AF2). The Rosetta filters weren’t used to filter ProteinMPNN designs because Rosetta models of ProteinMPNN outputs didn’t exist.

**Figure S1.**
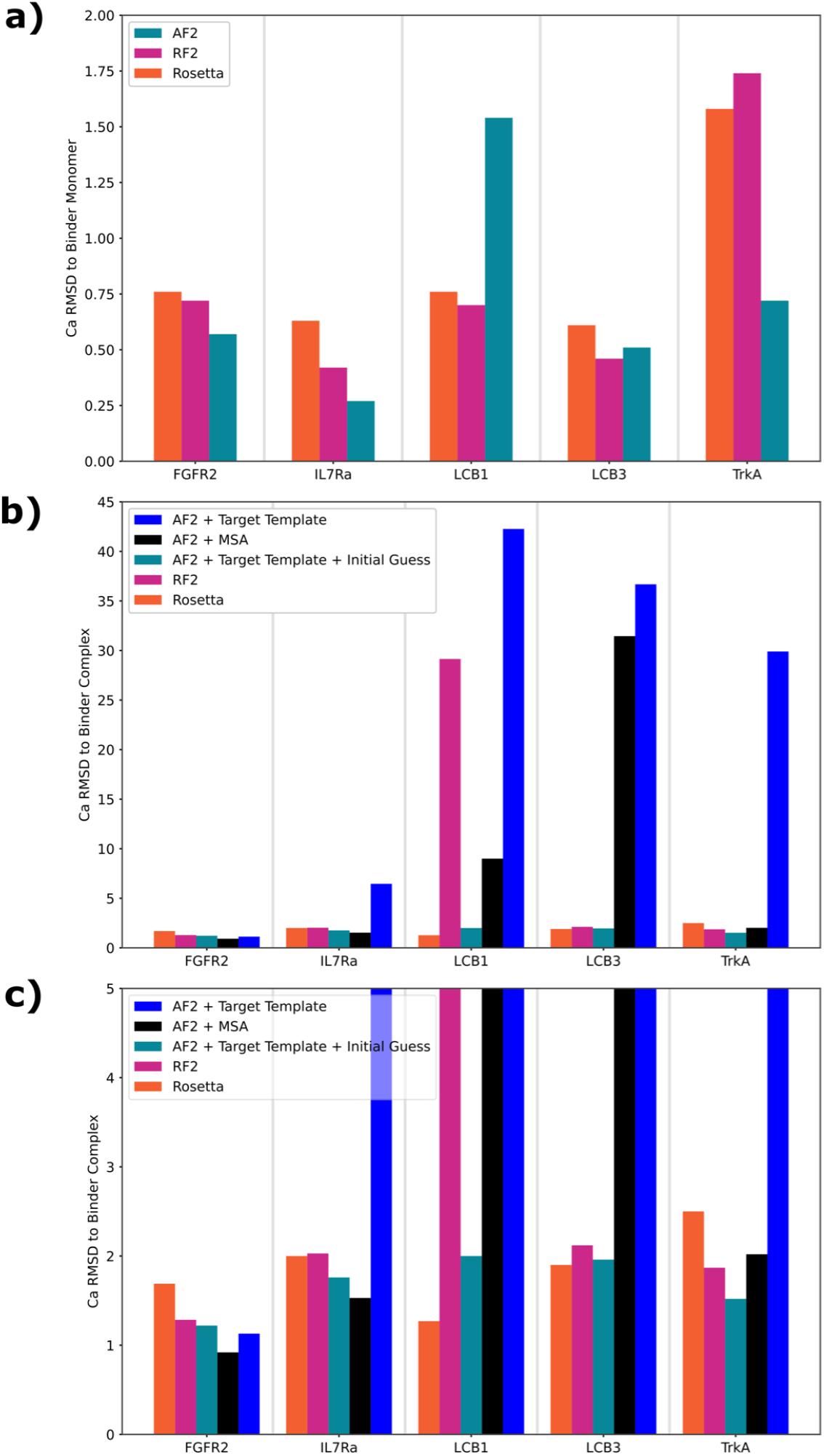
AF2 Prediction of Minibinder Crystal Structures. (A) The accuracy in Cα RMSD of AF2 and RF2 predictions of binder monomer structure for the five minibinder complex structures reported in Cao et al. (B) The accuracy in Cα RMSD of AF2 and RF2 predictions of binder complex structure for the five minibinder complex structures reported in Cao et al. (C) A close-up view of B.

**Figure S2.**
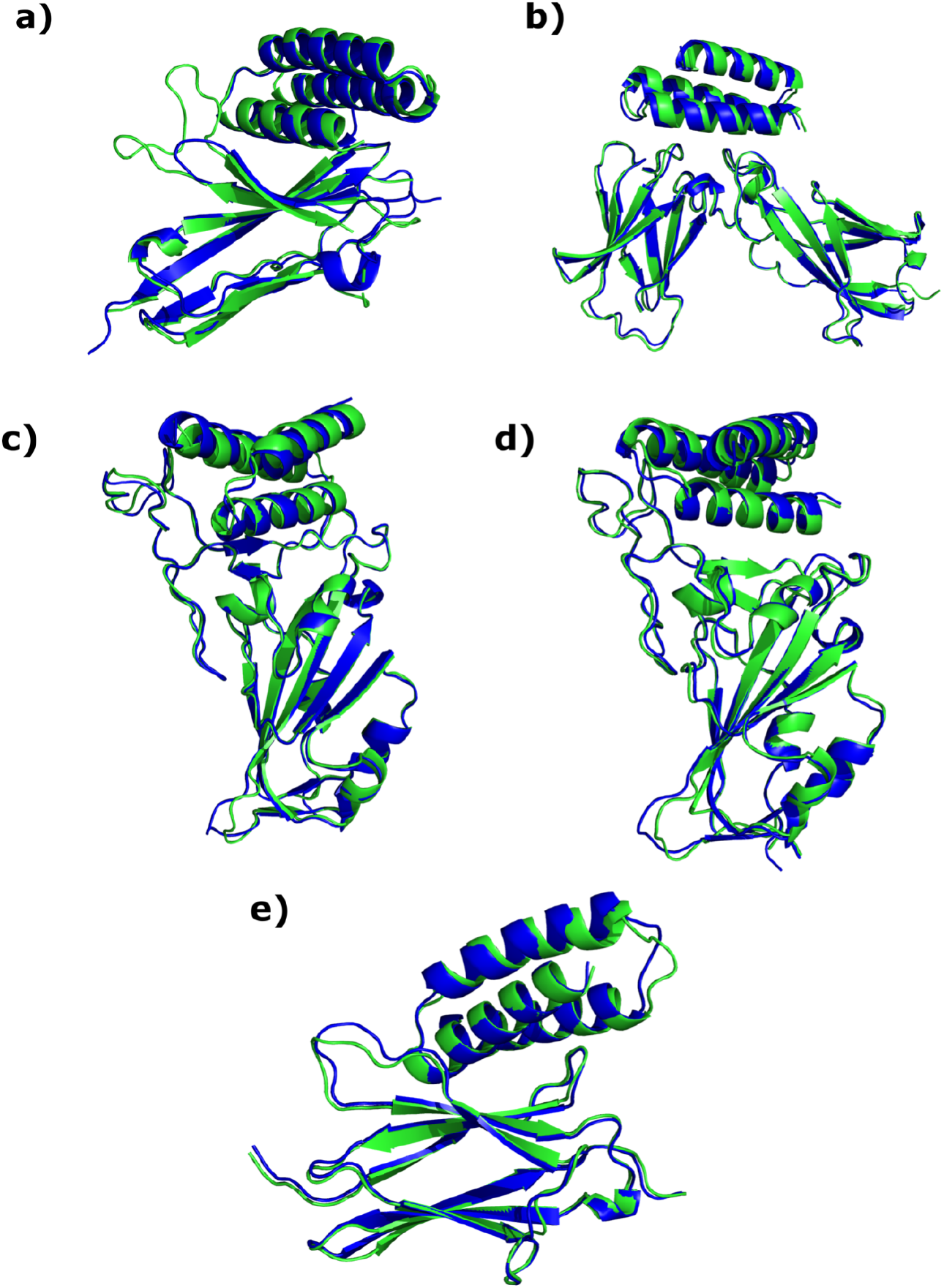
AF2 Complex Predictions (Blue) are close to the Crystal Structures (Green) All complex predictions shown were run using AF2 with target template and the Rosetta-design model as an initial guess. (A) FGFR2 (B) IL-7Rα (C) SARS-CoV-2 Spike (LCB1) (D) SARS-CoV-2 Spike (LCB3) (E) TrkA

**Figure S3.**
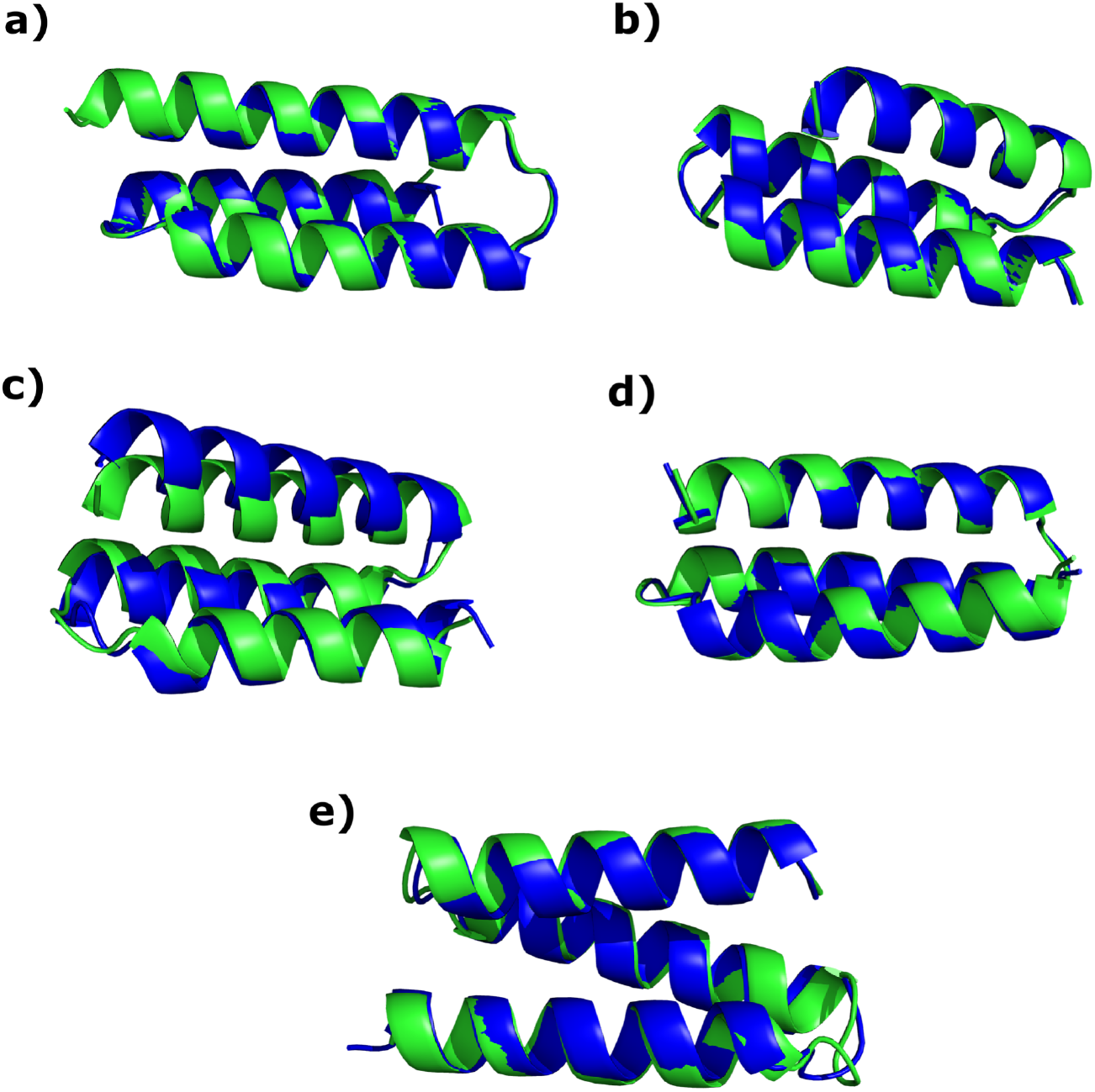
AF2 Monomer Predictions (Blue) are close to the Crystal Structures (Green) (A) FGFR2 (B) IL-7Rα (C) SARS-CoV-2 Spike (LCB1) (D) SARS-CoV-2 Spike (LCB3) (E) TrkA

**Figure S4.**
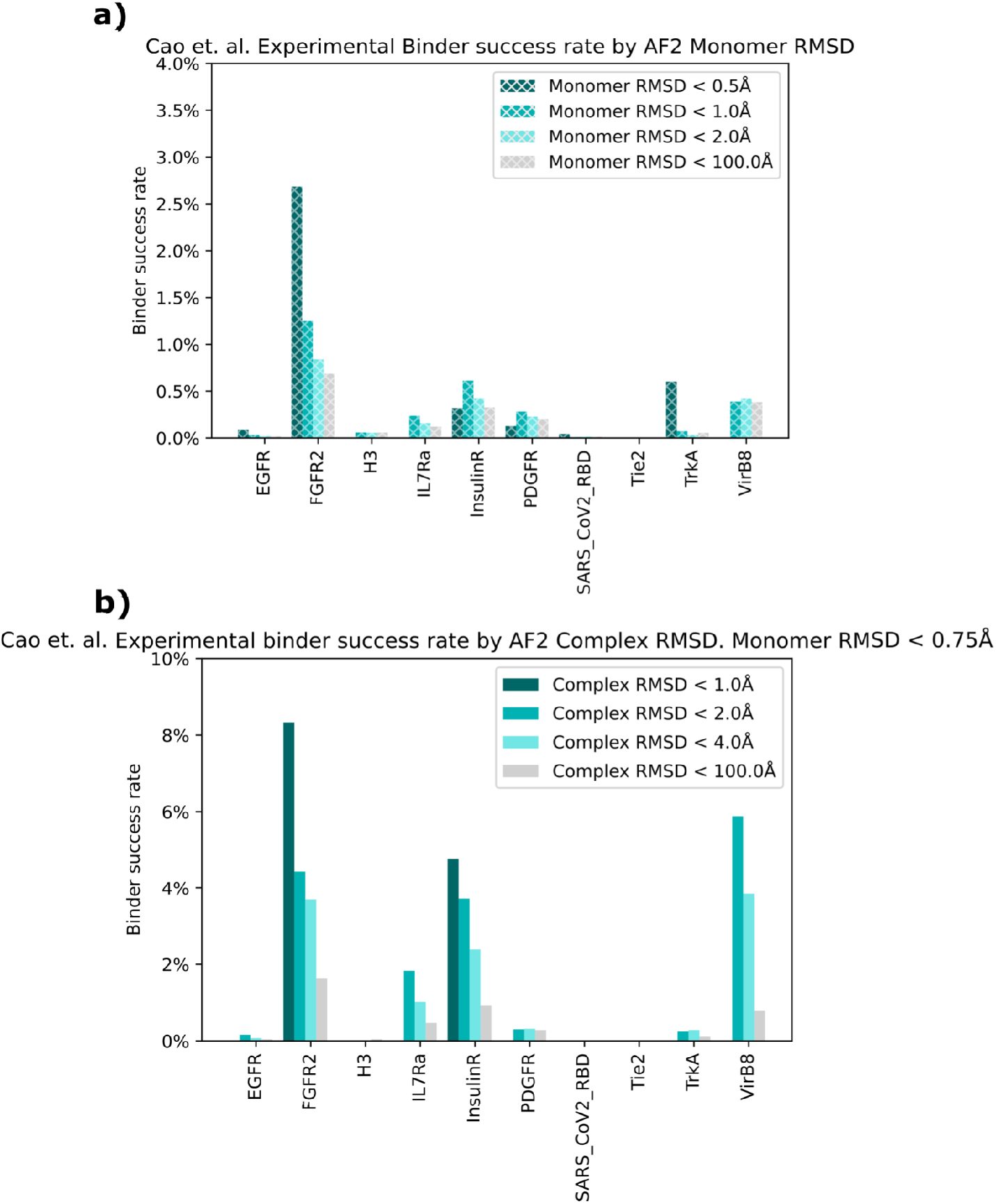
Retrospective Analysis of RMSD Metrics. (A) The retrospective experimental success rate (YSD SC_50_ < 4μM) for designs passing 4 thresholds of the RMSD difference between the Rosetta monomer structure and the AF2 monomer structure. (B) The retrospective experimental success rate (YSD SC_50_ < 4μM) for designs passing 4 thresholds of the RMSD difference between the Rosetta complex structure and the AF2 complex structure.

**Figure S5.**
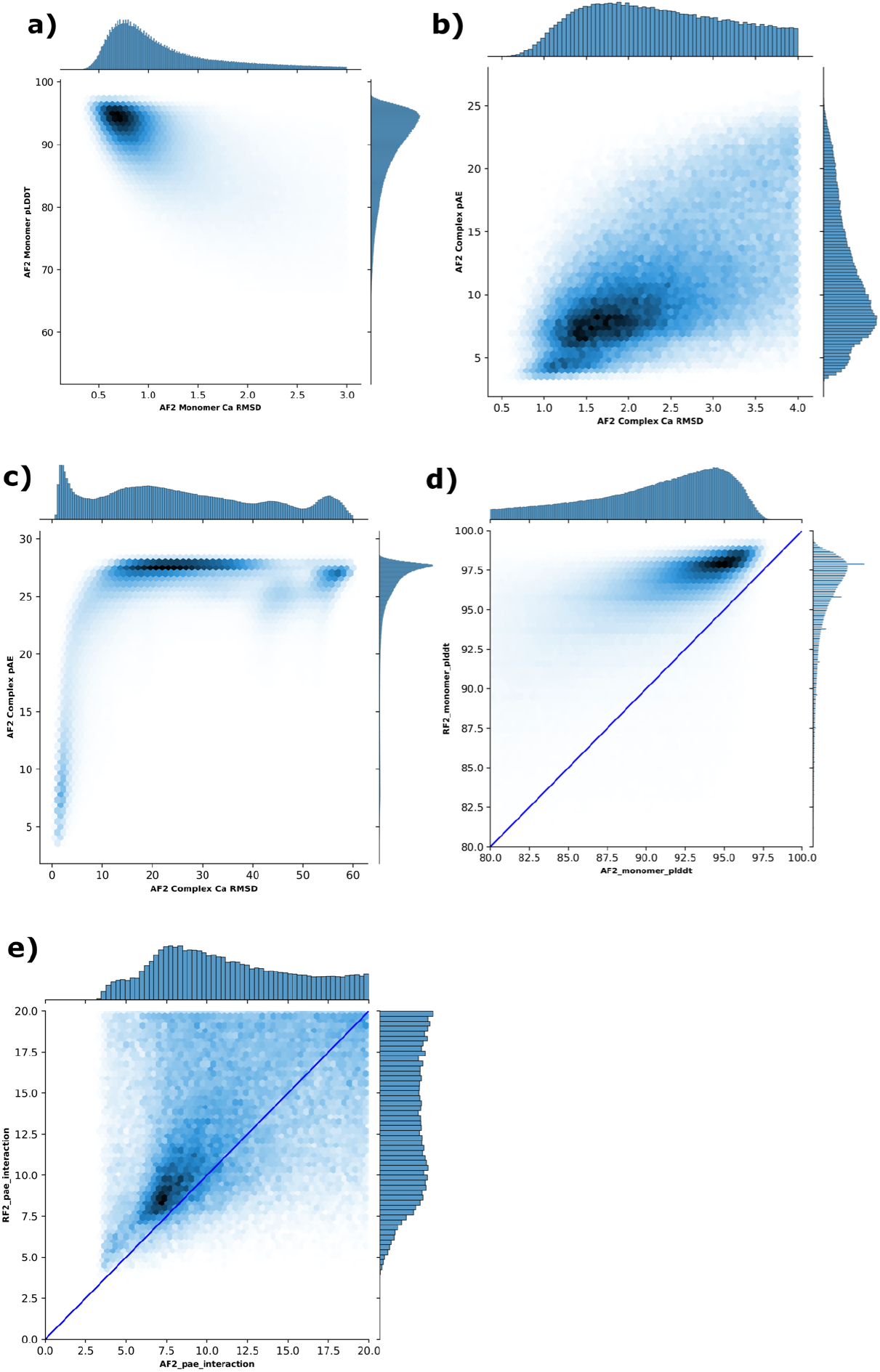
Correlation Between AF2 Confidence Metrics and AF2 RMSD. (A) For all binders in the dataset from Cao et al. the AF2 monomer confidence (pLDDT) and the RMSD difference between the Rosetta binder structure and the AF2 binder structure is plotted. (B) For all binders in the dataset from Cao et al. the AF2 complex confidence (pAE_interaction) and the RMSD difference between the Rosetta complex structure and the AF2 complex structure is plotted. (C) Zoomed-out version of B. (D) For all binders in the dataset from Cao et al. the AF2 monomer confidence (AF2_monomer_plddt) and the RF2 monomer confidence (RF2_monomer_plddt) is plotted. The line x=y is plotted in solid blue. (E) For all binders in the dataset from Cao et al. the AF2 complex confidence (AF2_pAE_interaction) and the RF2 complex confidence (RF2_pAE_interaction) is plotted. The line x=y is plotted in solid blue.

**Figure S6.**
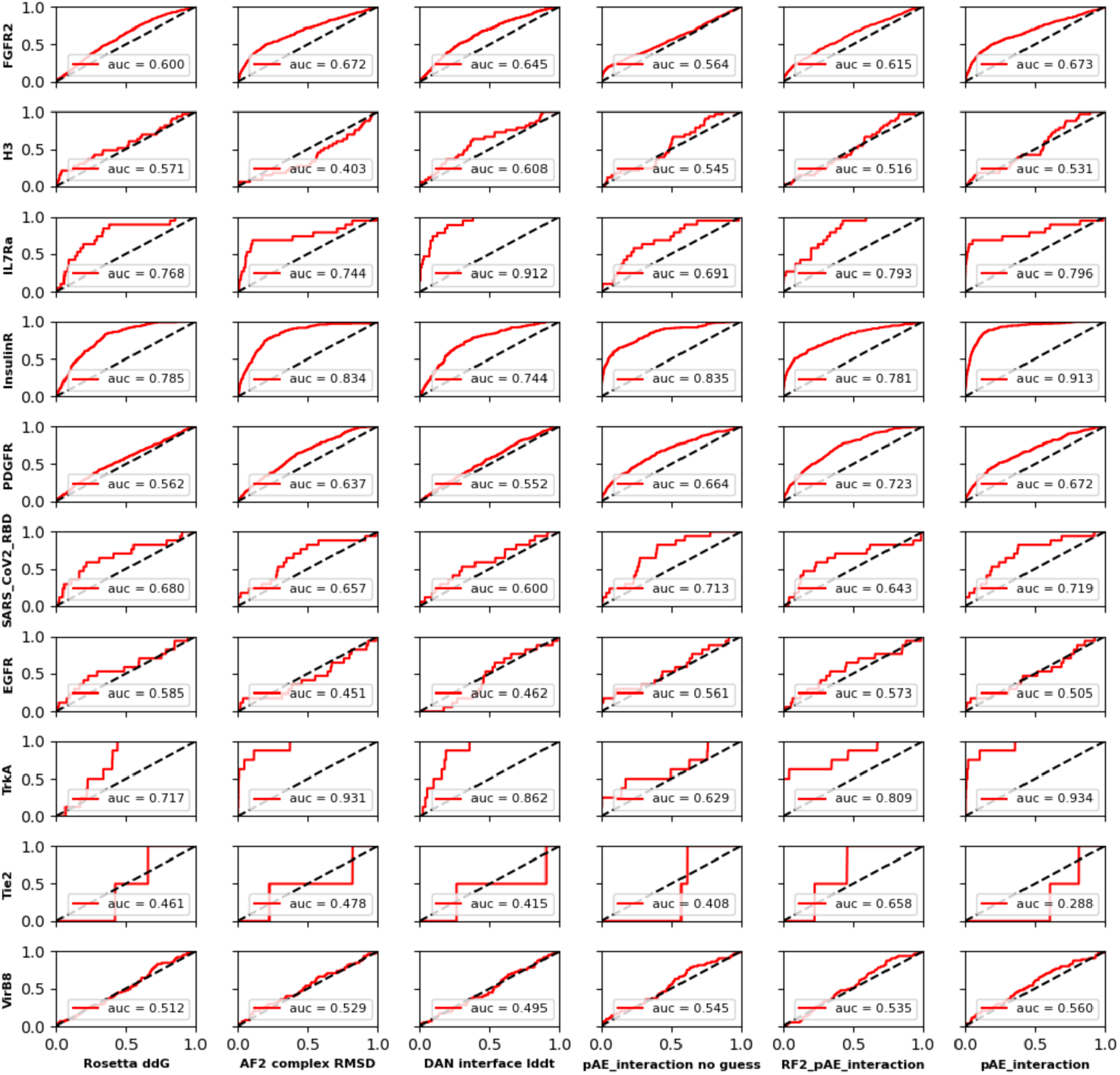
Retrospective Analysis of Complex Metrics. For each combination of 10 targets and 6 interface metrics a Receiver-Operator Characteristic curve quantifying the discriminatory power of the metric to separate successful designs (SC_50_ < 4 μM) from unsuccessful designs (SC_50_ >= 4 μM) is plotted. The True Positive Rate is plotted on the y-axis and the False Positive Rate on the x-axis. The Area Under the Curve (AUC) is included for each plot.

**Figure S7.**
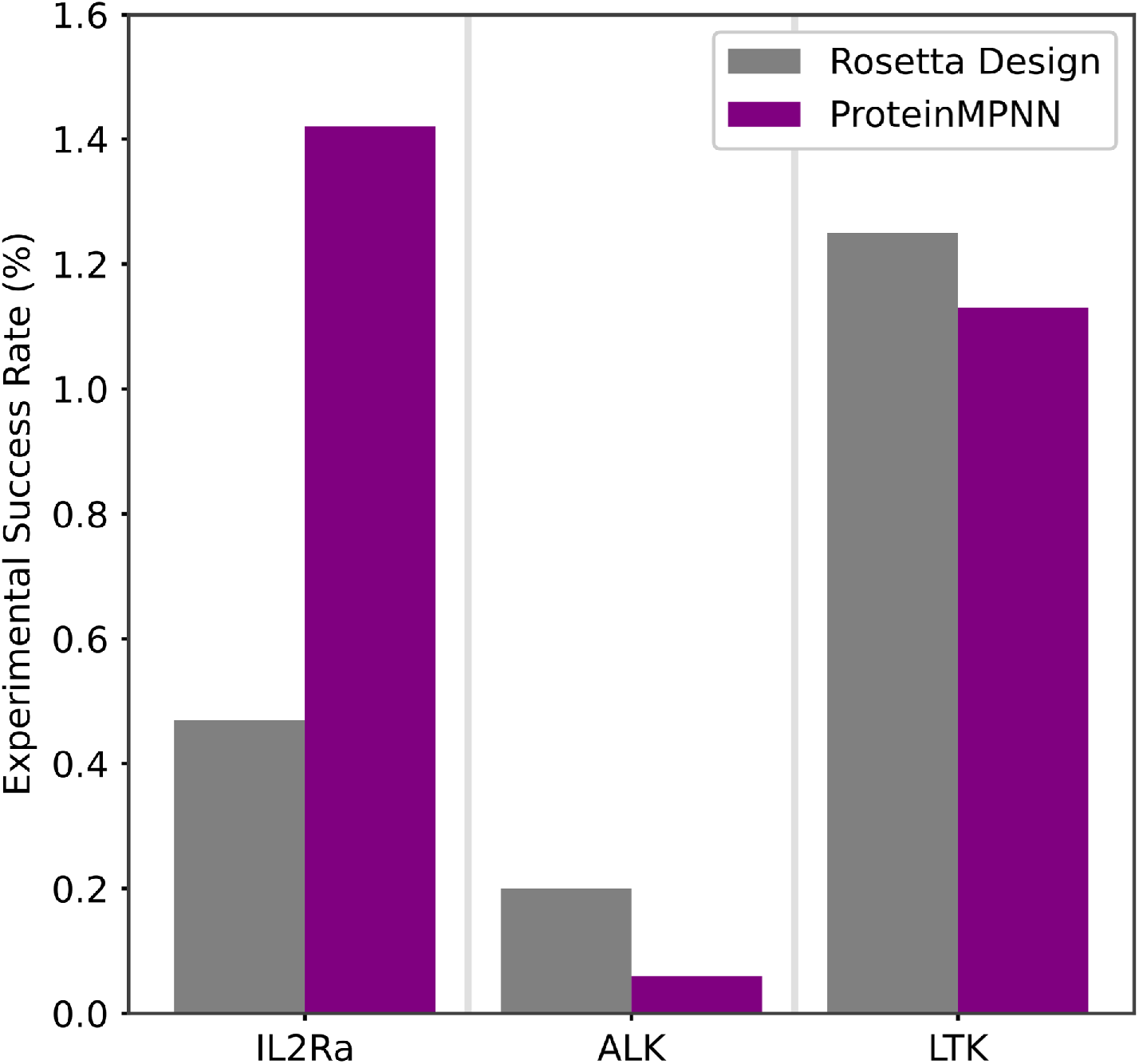
Experimental Success Rate of Binders Generated by ProteinMPNN versus Rosetta Design. The experimental success rate for libraries generated by redesign with FastDesign versus ProteinMPNN and filtered by DL-based filtering for three prospective targets. Neither redesign technique produced binders to IL10Rα or Site 2 of IL2Rα so these targets are not included here.

**Figure S8.**
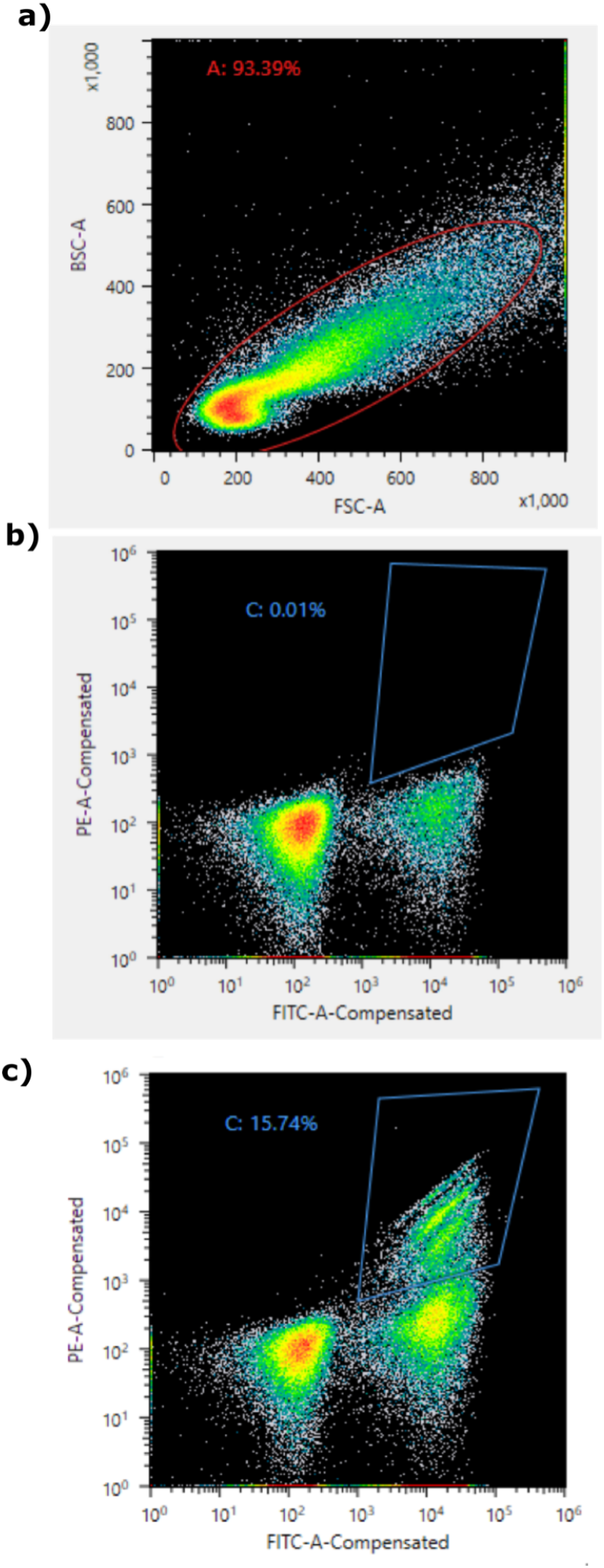
Gating Strategy for FACS Sorting of ALK Binder Library. a) Forward scatter (FSC) versus back scatter (BSC) plot with gate used in red. b) Negative sort with gate shown in blue. c) Binding sort with gate (same gate as used in b) shown in blue.

